# The EGFR inhibitor osimertinib promotes a dynamic Drug Tolerant Persister state marked by replication defects and genome instability

**DOI:** 10.64898/2026.06.19.733326

**Authors:** Kieron May, Giuditta Illuzzi, Matthew Martin, Jonathan Houseley

## Abstract

Osimertinib is the current standard-of-care for treatment-naive patients with EGFR mutation-positive advanced/metastatic non-small cell lung cancer (NSCLC), however resistance inevitably emerges. Osimertinib does not eradicate all cancer cells even in culture, leaving a long-lasting sub-population of Drug Tolerant Persister (DTP) cells that is common to many chemotherapeutics. The DTP population is non-proliferative and seemingly dormant, but resistant clones eventually emerge from the DTP state. Here we show that extensive DNA replication occurs in the DTP state and cells frequently progress through the cell cycle, though cell death is also frequent, such that cell division and cell loss are balanced and the population remains approximately constant. Cell cycling occurs with aberrant gene expression and abnormal DNA replication pattern, leading to DNA damage, extrachromosomal circular DNA formation and mitotic defects, such that replicating DTP cells are hypersensitive to low doses of ATM and ATR inhibitors. Our findings suggest that the DTP state is highly mutagenic and that targeting DNA repair in DTP cells has the potential to prevent the emergence of resistance through *de novo* mutations.

## Introduction

Non-small cell lung cancer (NSCLC) harbouring activating mutations in the epidermal growth factor receptor (EGFR) is highly responsive to EGFR-targeted therapies. Osimertinib (TAGRISSO®), a third-generation, irreversible EGFR tyrosine kinase inhibitor (TKI), selectively binds mutant EGFR (exon 19 deletions or L858R, with our without T790M) while sparing wild-type EGFR, reducing off-target toxicity (1). By occupying the ATP-binding pocket, osimertinib suppresses downstream signalling through the RAS–RAF–MEK–ERK and PI3K–AKT–mTOR pathways, leading to cyclin D1 downregulation, G1 cell-cycle arrest, and induction of mitochondrial apoptosis via pro-apoptotic proteins such as BIM (2–5). Reflecting this efficacy, osimertinib is recommended by the NCCN (US), ASCO (US), NICE (UK), and ESMO (Europe) as first-line therapy for advanced/metastatic EGFR-mutant NSCLC, adjuvant therapy following complete resection, and second-line therapy for patients with acquired T790M following prior EGFR TKI treatment (6–12).

Despite these advances, resistance to osimertinib inevitably emerges through diverse mechanisms, including secondary mutations or copy number variations in EGFR and other oncogenic drivers, oncogenic fusions, and lineage transformations such as small-cell lung cancer (13,14). Emerging evidence implicates extrachromosomal circular DNA (eccDNA) as a potential contributor to adaptive resistance (14,15). Notably, in EGFR-mutant NSCLC patients treated with osimertinib, plasma eccDNA sequencing revealed molecules overlapping the EGFR gene, with higher levels during treatment correlating with shorter progression-free and overall survival, supporting a role for eccDNA in treatment failure (15).

Resistance to targeted therapies can arise through pre-existing (intrinsic) mechanisms, driven by genetically or phenotypically heterogeneous tumour subclones present prior to therapy, or through acquired mechanisms, which develop during treatment via secondary mutations, copy number alterations, epigenetic reprogramming, lineage plasticity, or bypass pathway activation (16–21). A key intermediate state in this process is represented by drug-tolerant persister cells (DTPs): rare, phenotypically plastic subpopulations that survive therapy without classical resistance-conferring mutations. DTPs often enter a reversible slow-cycling or quiescent state maintained by epigenetic adaptations, transcriptional and metabolic rewiring, and alternative survival signalling (19,22,23). Upon drug withdrawal, DTPs can resume proliferation and regain drug sensitivity, but they also serve as a reservoir from which stable acquired resistance and relapse may emerge if additional genetic changes accumulate (23–28).

Consistent with this concept, we previously demonstrated that selumetinib (MEKi)-induced DTPs in colorectal cancer continue to replicate DNA despite exhibiting an aberrant transcriptional program (28). DNA replication under such compromised conditions can generate replication stress, which arises when replication forks stall due to oncogene-driven hyperproliferation, nucleotide depletion, limited replication machinery, DNA damage, or transcription–replication conflicts, ultimately promoting genomic instability (29–33). Targeted therapies can exacerbate this stress by perturbing cell-cycle control and replication dynamics, forcing surviving cells to replicate under suboptimal conditions (29). Error-prone fork restart, increased reliance on mutagenic repair pathways (e.g., translesion synthesis and microhomology-mediated end joining), and incomplete resolution of under-replicated DNA can produce de novo mutations and structural variants, which may restore pathway signalling or activate bypass mechanisms, thereby enabling acquired resistance to therapy (34–36).

Here, we investigate DNA replication defects and associated DNA repair events in DTPs maintained in a non-proliferative state by osimertinib.

## Materials and Methods

### Cell Lines and Culture Conditions

HCC4006, and NCI-H1975 human non–small cell lung cancer (NSCLC) cell lines were obtained from the American Type Culture Collection (ATCC) while PC9 NSCLC cells were obtained from Accegen. All lines were cultured in RPMI-1640 medium supplemented with GlutaMAX (Gibco, 61870-010), 10% fetal bovine serum (Gibco, A5256701), and 100 IU/mL penicillin–streptomycin (Sigma-Aldrich, P4333-100ML). Cells were maintained at 37°C in a humidified incubator with 5% CO□ and passaged at 70–80% confluence using 0.25% trypsin–EDTA (Gibco, 25200056). Cells were thawed from authenticated master stocks and used within 10–12 passages. Mycoplasma contamination was excluded by routine PCR-based testing. All experiments were performed using cells in logarithmic growth phase unless otherwise stated.

NSCLC Organoids were embedded in BME2 (Bio-techne, 33050-038) and overlaid with Advanced DMEM/F12 (Sigma)–based organoid medium supplemented with 1× GlutaMAX (Invitrogen, 12634-034) and 1× B27 supplement (Gibco, 17504001). The basal medium was further supplemented with N-acetylcysteine (1.25 mM), Wnt surrogate–Fc fusion protein (5 nM), A83-01 (500 nM), SB202190 (5 µM), Y-27632 (10 µM), and nicotinamide (2.5 mM). For long-term organoid expansion, media was additionally supplemented with recombinant human R-spondin-1 (final concentration 556 ng/mL), recombinant human Noggin (100 ng/mL), and recombinant human EGF (50 ng/mL). Medium was changed every 2–3 days and organoids were passaged by mechanical disruption upon reaching appropriate density. Organoids were maintained at 37°C in a humidified incubator with 5% CO□.

For long-term treatment experiments, cells were continuously exposed to 500 nM osimertinib dissolved in DMSO or equivalent vehicle control. Media containing drug or vehicle was replaced every 3–4 days. Final DMSO concentration did not exceed 0.1% (v/v) in any condition. Biological replicates represent independent cell cultures seeded using populations cultured independently for at least 5 passages prior. Unless otherwise specified, all experiments were performed using a minimum of two independent biological replicates.

### Generation of Stable FUCCI4 Reporter Lines

To enable cell cycle–resolved analyses, PC9 cells were engineered to stably express the FUCCI4 reporter (Addgene #83841) containing mKO2-CDT1 or Clover-GMNN marker proteins. Lentiviral particles were generated in HEK293T packaging cells via co-transfection of FUCCI4 with psPAX2 (Addgene #12260) and CMV-VSV-G (Addgene #8454) using Lipofectamine 2000 (Invitrogen, 11668019). Viral supernatants were collected 48 hours post-transfection, filtered (0.45 μm), supplemented with 3 μg/mL polybrene (Sigma-Aldrich, TR-1003-G), and applied to PC9 cells for 24 hours. Media was replaced after infection to minimize polybrene-associated toxicity. Transduced populations were enriched by fluorescence-activated cell sorting (FACS) of cells positive for either mKO2-CDT1 or Clover-GMNN marker proteins. Single cells were deposited into 96-well plates and clonally expanded. Clones were screened by live-cell fluorescence microscopy to confirm expected oscillatory expression patterns consistent with G1, S, and G2/M transitions and to exclude clones exhibiting aberrant cell cycle timing or reporter instability. Only clones demonstrating stable reporter dynamics over multiple passages were used for downstream experiments. Reporter stability was confirmed for at least 10 passages prior to experimental deployment.

### Drug Treatment Regimens

For adaptation and persistence models, cells were continuously treated with 500 nM osimertinib for the stated duration up to a maximum of 58 days. PC9 osimertinib resistant lines were generated by seeding cells and continually exposing them to 500 nM osimertinib until the culture vessel had reached 100% confluence. Resistance was confirmed by re-seeding and confirming similar growth kinetics in untreated, 500nM and 1µM osimertinib conditions. Dose–response assays were performed using a half-log dilution series ranging from 10 μM to 1 nM, treating for 4 days. Each condition was performed in technical triplicate. Vehicle controls were included in all experiments to control for DMSO exposure. Additional DNA damage response inhibitors were used at the following concentrations: Olaparib (300 nM), Saruparib (30 nM), Ceralasertib (100 nM), AZD7648 (100 nM), RP-3500 (100 nM), AZD1390 (30 nM), AZD1775 (50 nM), and AZD4641 (200 nM). All small-molecule inhibitors were synthesized according to published methods.

### TrAEL-seq

TrAEL-seq was performed to map replication-associated DNA 3′ ends genome-wide as previously described (37,38).

Approximately 1 × 10□ cells per sample were embedded in CleanCut agarose (Bio-Rad, 1703594) and lysed overnight at 50°C in mammalian digestion buffer (0.5 M EDTA pH 8.0, 1% sodium lauroyl sarcosinate) supplemented with 1 mg/mL proteinase K to ensure complete protein removal while preserving DNA integrity. Plugs were washed extensively in TE buffer and treated with RNase T1 to remove residual RNA. DNA 3′ ends were A-tailed using Terminal Transferase (NEB, M0315L) in the presence of ATP and ligated to biotinylated TrAEL-seq adaptors using T4 RNA

Ligase 2 truncated KQ (NEB, M0373L). Following β-agarase digestion (NEB, M0392S), DNA was ethanol precipitated and extended using Bst 2.0 WarmStart DNA Polymerase (NEB, M0538S) to generate double-stranded products compatible with library preparation. DNA was fragmented to 200–400 bp using a Covaris E220 system under standardized acoustic parameters. Biotinylated fragments were enriched using Dynabeads MyOne Streptavidin C1 (Thermo Fisher, 65001). Libraries were prepared using the NEBNext Ultra II DNA Library Prep Kit (NEB, E7645S) with USER enzyme treatment (NEB, M5505S). PCR amplification cycles (typically 8–12) were optimized using test amplifications to avoid over-amplification bias. Libraries were purified using AMPure XP beads, quality assessed by Agilent Bioanalyzer, quantified by KAPA Library Quantification Kit, pooled equimolarly, and sequenced (75 bp single-end) on an Illumina NextSeq 500 to a depth of ∼10 million reads per sample. Independent biological replicates were generated from separate cultures.

### RNA Sequencing of flow-sorted cells

Cells were fixed with glyoxal as described to preserve RNA integrity during sorting (39). Following fixation cells were saponin-permeabilized and using FACS were sorted into G1, S, and G2 populations. Cells were first gated based on DAPI content, followed by refinement using mKO2-CDT1 and Clover-GMNN marker expression, and were collected directly into TRI Reagent for RNA extraction.

Total RNA was extracted using TRI Reagent (Sigma, T9424) followed by organic phase separation and isopropanol precipitation. RNA purity was assessed by spectrophotometry. Poly(A)+ RNA was isolated using the NEBNext Poly(A) mRNA Magnetic Isolation Module (NEB, E7490S). Strand-specific libraries were prepared using the NEBNext Ultra II Directional RNA Library Prep Kit (NEB, E7760S). Libraries were amplified for 10–18 PCR cycles based on test amplifications carried out on a small portion of the sample. Following AMPure XP bead purification, libraries were quality assessed by Agilent Bioanalyzer, quantified by KAPA Library Quantification Kit. Sequencing was performed (75 bp single-end) to an average depth of ∼24 million reads per sample.

### Circle-seq

Circle-seq was developed to enable sensitive profiling of extrachromosomal circular DNA (eccDNA) from low-input human cell populations. The method is based on selective enzymatic depletion of linear genomic DNA followed by unbiased amplification of circular DNA templates and high-throughput sequencing. Approximately 10,000 cells per sample were lysed in 10 µL Buffer RLT+ (Qiagen, 1053393) and incubated at room temperature for 20 minutes, followed by DNA purification using 6 µL AMPure XP beads and elution in 5 µL 0.1× TE. Purified material was then treated with a protein digestion mix (0.75 µL 10× NEB4 buffer (New England Biolabs, B7004), 0.1125 µL 10% Triton X-100 (Sigma-Aldrich, X100-100ML), 1.4375 µL nuclease-free water, and 0.2 µL thermolabile proteinase K (NEB, P8111S)) and incubated at 37°C for 3 hours followed by heat inactivation at 85°C for 10 minutes.

In some reactions, linear genomic DNA was depleted by sequential digestion using RecBCD and Exonuclease I under conditions favouring linear DNA degradation. Specifically, RecBCD exonuclease (NEB, M0345S), which preferentially degrades linear double-stranded DNA, and Exonuclease I (NEB, M0293L), which degrades single-stranded linear DNA species, were applied to remove chromosomal DNA fragments. Exonuclease digestion was performed by adding 2.5 µL of digestion mix (0.25 µL 10× NEB4 buffer (New England Biolabs, B7004), 0.125 µL 100 mM ATP, 0.2 µL RecBCD, 0.1 µL Exonuclease I, and 1.825 µL nuclease-free water) and incubating at 37°C overnight. This digestion was repeated for a total of four overnight incubations with incremental ATP supplementation (0.025 µL per cycle) prior to heat inactivation at 80°C for 20 minutes. Efficiency of linear DNA removal was verified by qPCR targeting mitochondrial DNA and rDNA loci. Samples demonstrating insufficient depletion were excluded, with a depletion efficiency threshold defined as ≥5 Cq cycles for a primer pair targeting a high-copy chromosomal locus (e.g. rDNA) relative to undigested input controls. Following exonuclease treatment, DNA was purified using a 1:1 ratio of AMPure XP beads and eluted in 7 µL 0.1× TE. Circular DNA was amplified using the REPLI-g Mini Kit (Qiagen, 150023). Multiple displacement amplification (MDA) was selected due to the strand-displacing activity of phi29 DNA polymerase, which enables rolling-circle amplification of circular templates with high fidelity. Amplification reactions were initiated by adding 0.44 µL Buffer DLB and incubating at room temperature for 3 minutes, followed by addition of 0.6 µL stop solution and 12 µL master mix (11.6 µL REPLI-g buffer and 0.4 µL enzyme). Reactions were performed under isothermal conditions (30°C, ∼16 hours), followed by enzyme inactivation at 65°C for 20 minutes, and DNA purification using 0.8× AMPure XP beads with elution in 15 µL 0.1× TE.

To minimise amplification bias, reactions were limited to the minimum input required to generate sufficient material for sequencing. In addition, parallel technical replicates were performed to assess reproducibility. Amplified DNA was purified using AMPure XP beads and quantified using Qubit dsDNA HS assay. Samples were diluted to 200 pg/µL prior to library preparation. Sequencing libraries were prepared using the Nextera XT DNA library preparation kit (Illumina, FC-131-1024). For tagmentation, 1 µL of diluted DNA was added to 2 µL tagment DNA buffer, incubated on ice for 2 minutes, followed by addition of 1 µL Amplicon Tagment Mix (ATM). Samples were incubated at 55°C for 5 minutes, neutralised with 1 µL NT buffer (5 minutes, room temperature), and amplified by PCR using 1 µL each of index primers (5 µM) and 3 µL NPM under the following conditions: 72°C for 3 minutes, 95°C for 30 seconds, 12 cycles of (95°C for 10 seconds, 55°C for 30 seconds, 75°C for 30 seconds), and a final extension at 72°C for 5 minutes. Libraries were purified using 12 µL AMPure XP beads and eluted in 10.5 µL 0.1× TE. Sequencing was performed (75 bp paired-end) on an AVITI platform (∼10 million reads/sample). Paired-end sequencing was selected to improve mapping confidence across circular junction regions and repetitive genomic elements.

### Bioinformatic Analysis

All sequencing datasets were processed by the Babraham Bioinformatics Facility using standardized pipelines. Reads were aligned to GRCh38 supplemented with rDNA sequence (U13369.1).

TrAEL-seq reads were adapter-trimmed using Trim Galore (v0.6.5) and aligned using Bowtie2 (v2.4.1). UMI-based deduplication was performed to remove PCR duplicates. Only uniquely mapped reads (MAPQ ≥ 20) were retained. Replication initiation zones were identified using OKseqHMM (threshold = 8; bin size = 10 kb; smoothing window = 25 bins) (40). Strand bias metrics were computed in SeqMonk (v1.48.1). R scripts were used to calculate replication fork directionality, distance-to-origin distributions, and aggregate metaplots (37).

For RNA-seq, trimmed reads were aligned with HISAT2 (v2.1.0). Gene-level counts preformed in SeqMonk (v1.48.1) were imported into DESeq2 for differential expression analysis (Benjamini–Hochberg FDR < 0.05). Principal component analysis confirmed replicate clustering by condition. Gene ontology enrichment was performed using Gorilla (41).

For Circle-seq, trimmed reads were aligned using Bowtie2 (v2.4.1). Circular DNA regions were identified by quantifying read density in 500 bp bins across the genome using SeqMonk (v1.48.1). Bins exceeding defined enrichment thresholds relative to background were retained and adjacent bins merged. Circular junction-supporting read pairs were examined using CircleFinder (v2.0.0.1) to confirm continuity consistent with circular templates.

Data is available from GEO accession number: GSE335848

### EdU Incorporation and Immunofluorescence

For DNA replication assays, cells were incubated with 1 µM EdU for 2 hours prior to fixation in 4% paraformaldehyde. Following permeabilization (0.5% Triton X-100), EdU was detected using Alexa Fluor 647 azide (Lumiprobe, A6830) via copper-catalyzed click chemistry. For DNA damage analyses, cells were labelled with 10 μM EdU for 30 minutes and stained with antibodies against γH2AX (Upstate, JBW301), phospho-RPA (Thermo, A300-245A), or phospho-KAP1 (Thermo, A300-767A). Images were acquired using Nikon Eclipse Ti2 or Olympus ScanR automated microscopy systems under fixed acquisition settings. Quantification was performed using CellProfiler (v4.2.4) or Olympus AI segmentation pipelines. ≥500 cells per condition per replicate were analysed.

### Cell Viability Assays

Cells were seeded at 5,000 cells per well in 96-well plates and treated with serial drug dilutions. After 96 hours, CellTiter-Glo 2.0 reagent (Promega, G9241) was added and luminescence measured using a SpectraMax iD5 plate reader. Dose–response curves were fit using nonlinear regression in GraphPad Prism 10. IC[[values were derived from at least three independent biological replicates.

### Mitotic Chromosome Segregation Analysis

Cells treated ± 500nM osimertinib for 28 days were fixed and stained for phospho-Histone H3 Ser10 (Cell Signalling, 9701S). Mitotic cells were imaged using z-stack acquisition with 0.6 µM increments and scored in a blinded manner for segregation defects. At least 50 mitotic cells per condition per replicate were analysed.

### Live-Cell Imaging

Live imaging was performed using an Incucyte SX5 system. Confluence was quantified using AI-based segmentation algorithms. For cell fate analyses, individual FUCCI-positive cells were manually tracked for 24 hours and classified based on cell cycle progression and division events.

### Statistical Analysis

All statistical analyses were performed using GraphPad Prism 10. Data are presented as mean ± SD unless otherwise specified. Statistical tests are given in figure legends.

## Results

### Extensive DNA replication occurs in the DTP state

Our previous finding that cells in the DTP state can enter the cell cycle led us to examine DNA replication in cells treated with osimertinib, the current standard of care for EGFR-mutant NSCLC. Single cell analysis has revealed ongoing proliferation in the first 14 days after osimertinib treatment (42), but it remains unclear if this persists through the DTP state or simply represents a tail of cells yet to succumb to cell cycle arrest. This is important as proliferative resistant populations emerge from the DTP state (23,26–28).

We treated PC9, a non-small cell lung carcinoma line carrying an activating exon 19 EGFR deletion mutation (E746-A750), with 500 nM osimertinib: this concentration robustly induces a DTP state (43,44) with the PC9 response to osimertinib being essentially uniform from 10 nM-1 µM (Figure S1A). After an initial wave of cell death lasting 1-2 days, treated cultures reproducibly formed a stable DTP state in which cells remained viable but did not require passaging for ∼3 months until resistant clones suddenly emerged (Figures 1A,B). To quantify replicating cells, cultures were pulsed for 2 h with 5-ethynyl-2’-deoxyuridine (EdU) prior to harvest after 0, 7, 14 and 28 days of osimertinib treatment, and EdU positive cells quantified as a fraction of the population by high-throughput imaging (Figure 1C). Remarkably, although populations were not proliferative as a whole, many EdU-positive cells were observed; absolute amounts across the first month of treatment varied between experiments likely due to small variations in seeding density (compare the day 7 and day 14 data between Figures 1B,E), but consistently stabilised at 35-40% which is ∼half the fraction of replicating cells in the untreated, rapidly proliferating population. This is far greater than we observed for selumetinib (MEKi) treated cells (28), but was reproduced in HCC4006 cells (Figure S1B,C), while in an NSCLC organoid with a heterozygous EGFR L858R mutation 50 nM osimertinib suppressed replication at 4 days, but the replicating population returned by Day 11 (Figure S1D), showing that the osimertinib-induced DTP state is unexpectedly dynamic.

**Figure 1:**
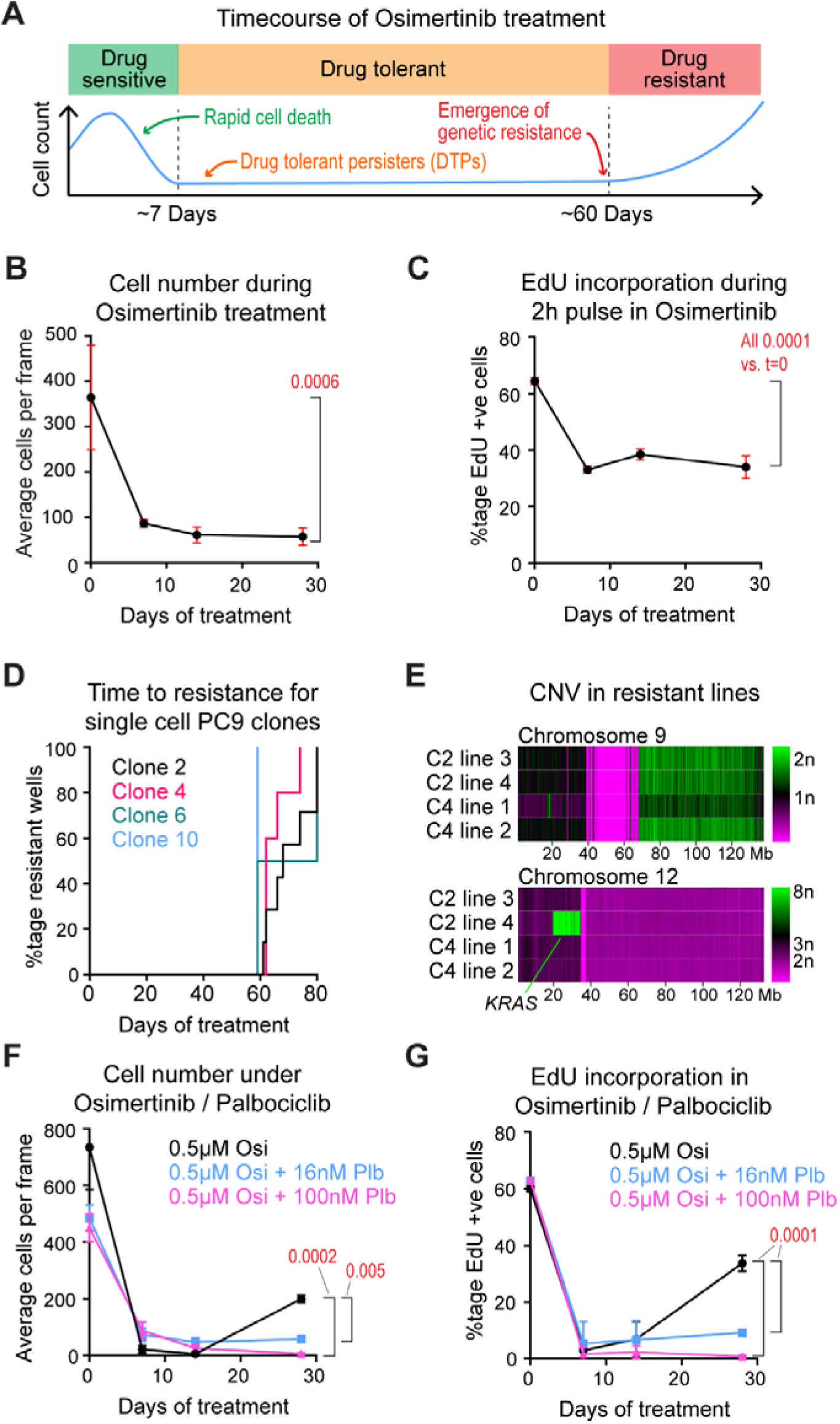
Extensive DNA replication occurs in the DTP state. A) Schematic illustrating population dynamics during osimertinib treatment. Cancer cells initially undergo a drug-sensitive phase characterised by rapid cell death, followed by the emergence of a small drug-tolerant population exhibiting relative dormancy (DTP state), prior to the development of genetically resistant clones. B) Quantification of PC9 cell numbers over 28 days of 500 nM osimertinib treatment. Statistical significance was determined by one-way ANOVA comparing all time points to day 0, with Dunnett’s multiple comparisons correction, n=3, error bars ±1 SD. C) Quantification of the proportion of PC9 cells staining positive after a 2 h EdU pulse, over 28 days of 500 nM osimertinib treatment. Statistical significance was determined by one-way ANOVA comparing all time points to day 0, with Dunnett’s multiple comparisons correction, n=3, error bars ±1 SD. D) Time required for single-cell-derived PC9 clones to acquire resistance to osimertinib. Resistance was defined as cultures reaching confluence under continuous 500 nM osimertinib treatment. 12 cultures of each clone were started, though some were lost to contamination, final n=5-7. E) Quantification of genomic copy number based on genome resequencing across chromosomes 9 and 12 in single-cell-derived PC9 clones following acquisition of osimertinib resistance. F) Quantification of PC9 cell numbers over 28 days of treatment with 500 nM osimertinib alone or in combination with 16 nM or 100 nM palbociclib. Statistical significance was determined by two-way ANOVA comparing combination treatments to single-agent osimertinib, with Dunnett’s multiple comparisons correction, n=3, error bars ±1 SD. G) Quantification of the proportion of PC9 cells staining positive for EdU (2-hour pulse, 1 µM) over 28 days of treatment with 500 nM osimertinib alone or in combination with 16 nM or 100 nM palbociclib. Statistical significance was determined by two-way ANOVA with Tukey’s multiple comparisons correction, n=3, error bars ±1 SD.

A defining characteristic of the DTP state is the potential to acquire resistance (19). Resistance can emerge though acquired mutations but also through selection of resistant alleles from a heterogeneous starting population. To minimize heterogeneity, four clonal populations of PC9 cells were generated from individual cells, all of which became resistant to 500 nM osimertinib after 2-3 months in the DTP state without passaging (Figure 1D). Resistance was scored based on cultures reaching confluence, and confirmed by measuring growth of resistant clones and single cell clones derived from those resistant clones (Figure S1E); in all cases the resistant lines were slightly slower proliferating than the parental but growth was unaffected by osimertinib. Genome re-sequencing of 2 resistant lines each derived from 2 single cell clones revealed large CNVs in one line from each clone, one involving an amplification of *KRAS*, but other resistant lines showed no detectable CNVs (Figure 1E). The emergence of resistance from clonal starting populations after an extended non-proliferative period is consistent with osimertinib resistance being acquired in the DTP state, as a resistance mutation in the starting population should render these immediately resistant to treatment.

Osimertinib causes G1 arrest by inhibiting EGFR and thereby preventing activation of the MEK-ERK pathway which promotes the G1-S transition through activation of cyclin D by CDK4/6 (45–47). We have previously shown that MEK inhibition can be strengthened in the DTP state using low concentrations of the CDK4/6 inhibitor palbociclib (28). 16 nM and 100 nM palbociclib, which have minimal effect on PC9 proliferation as a single agent (Figure S1F), dramatically reduced the proportion of cells undergoing DNA replication after 28 days treatment in combination with 500 nM osimertinib, and also dramatically reduced the number of cells (Figure 1E,F). In the presence of palbociclib, the DTP state was therefore no longer stable, and no resistant clones emerged simply due to all the cells losing viability in the first 30-60 days. This shows that continued proliferation under osimertinib treatment is required for the ongoing maintenance of the DTP state and the emergence of resistance.

These experiments reveal that the DTP state induced by osimertinib is not composed of long-lived, stable non-replicating cells that persist in drug but rather a population in which frequent DNA replication must be balanced by a similar rate of cell death.

### The DTP cell cycle involves frequent cell death and aberrant gene expression

Low-level EdU incorporation could occur through DNA repair rather than DNA replication, however EdU intensity is equivalent for osimertinib treated cells and control cells, showing that equivalent DNA synthesis occurs (Figure 2A). Furthermore, EdU positive cells become Cyclin B1 positive in late S/G2 (Figure 2B), and anaphase cells are only reduced by ∼50% compared to the untreated control (Figure 2C). The replicating cells may represent a small resistant sub-population, however on extended 3-day EdU exposure ∼70% of cells incorporated EdU even in 14-day treated cultures for which only ∼10% of cells incorporate EdU in 2 h (Figure S2), indicating that most DTP cells periodically enter the cell cycle, and cell cycle progression then follows normal dynamics.

**Figure 2:**
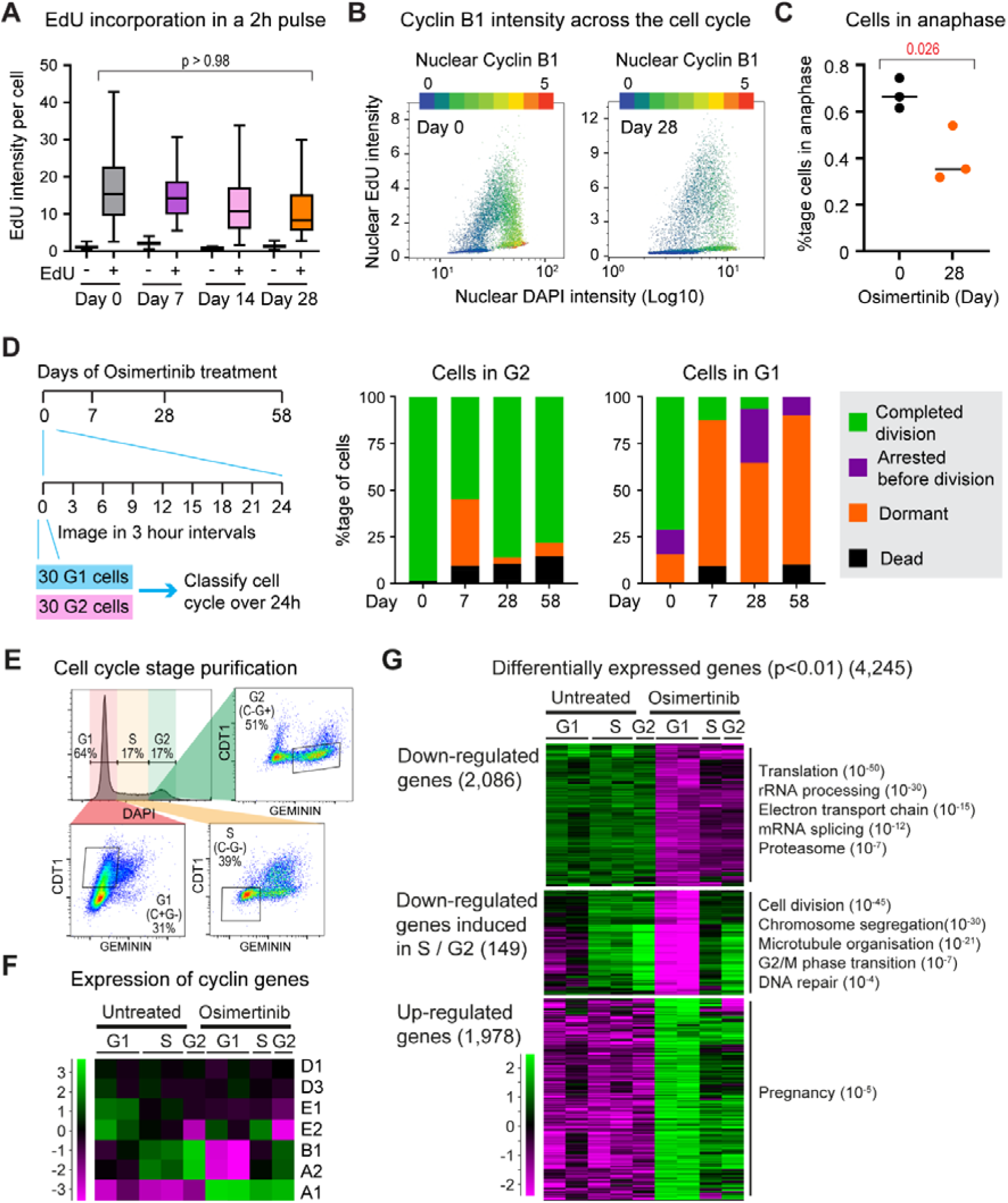
The DTP cell cycle involves frequent cell death and aberrant gene expression. A) Quantification of EdU intensity per cell in EdU-positive and EdU-negative populations after a 2 h EdU pulse at days 0–28 of 500 nM osimertinib treatment. Statistical significance was determined by repeated-measures one-way ANOVA comparing EdU-positive populations at days 0 and 28, with Bonferroni correction. B) Quantification of nuclear Cyclin B1 intensity per cell shown on a colour scale, relative to DAPI and EdU intensity after a 2 h EdU pulse at days 0–28 of 500 nM osimertinib treatment. C) Quantification of the proportion of cells undergoing anaphase at days 0–28 of 500 nM osimertinib treatment, identified by Histone H3S10 phosphorylation and chromosomal morphology on blinded samples. Statistical significance was determined by unpaired t-test, n=3. D) Schematic and quantification of live-cell imaging of 30 PC9 cells monitored over 24 hours (imaged every 3 hours) at days 0, 7, 28, and 58 of 500 nM osimertinib treatment. Cells were initially selected in either G1 or G2 cell cycle phases and their outcomes at the end of the observation period were categorised. E) Schematic of cell cycle sorting strategy using FACS, with initial gating based on DAPI intensity followed by refinement using CDT1(30–120) and GMNN(1–110) reporter expression. F) Heatmap showing log_2_ fold-change values of Cyclin genes determined in cell cycle-resolved PC9 populations following 0 and 28 days of 500 nM osimertinib treatment. G) Heatmap of log_2_ fold-change values for genes that are significantly downregulated, upregulated, or differentially regulated across the cell cycle (e.g. downregulated in G1 but induced in S/G2) in PC9 cells following 28 days of osimertinib treatment relative to untreated controls, with associated enriched gene ontologies.

To monitor the cell cycle during osimertinib treatment, we generated a cell-cycle reporter Fucci4 PC9 line containing CDT1(30–120) and GMNN(1–110) reporter proteins for tracking of G1 and G2 phases respectively (48). Fucci PC9 cells were treated with 500 nM osimertinib for 0, 7, 28 or 58 days then imaged every 3 h for 24 h using an Incucyte SX5. 30 G1 and 30 G2 cells were selected at random at each time point and their fate across 24 h determined (Figure 2D): G2 cells in the untreated population almost all successfully divided, but with time in the DTP state increasing numbers of G2 cells died either before or after mitosis, with others arresting without dividing. This provides an insight into why the population remains stable despite ongoing high levels of DNA replication. Curiously, while G1 cells marked by Cdt1 in the untreated population divided within 24 h, Cdt1-positive cells in the osimertinib treated populations did not, despite the fact that EdU assays show cells frequently enter S phase. We suspect that cells scored as Cdt1-positive have been efficiently arrested in G1 by the drug, whereas cells which enter S phase do so with less accumulation of Cdt1 and are not scored as Cdt1-positive against background.

We then used the Fucci derivative combined with DAPI staining to sort cells in G1, S and G2 after glyoxal staining to recover high quality RNA (Figure 2E). This worked effectively in Day 0 cells, yielding two biological replicate libraries each for G1 and S, as well as one library of G2 cells (which were much rarer). After 28 days, there were few cells in the cultures and much debris, but we sorted sufficient cells for two biological replicate G1 libraries and one each of S and G2 cells. We examined cyclin mRNA levels to validate the sorting (Figure 2F): cyclin D mRNA (*CCND1* and *CCND3* only, *CCND2* was not detectably expressed) did not change greatly since cyclin D is primarily post-transcriptionally regulated; cyclin E mRNA (*CCNE1* and *CCNE2*) was strongly expressed in G1 in untreated cells but not in treated cells, coherent with expression being induced downstream of cyclin D-CDK4/6 which is blocked by osimertinib; cyclins A2 and B1 (*CCNA2* and *CCNB1*) increased progressively from G1 to S to G2 at both Day 0 and 28. Unexpectedly *CCNA1*, encoding a cyclin normally only expressed in testes and brain but which is induced in response to DNA damage, was highly expressed in osimertinib-treated cells (49–51).

4,245 genes changed significantly (p<0.01) in the G1 sets between Day 0 and Day 28 (Figure 2G): down-regulated genes were highly enriched for translation (ie: ribosomes and ribosome synthesis), mitochondrial proteins, splicing and proteosome components, all of which are critical for rapid proliferation. A small number (149 of 2,235) of these genes enriched for mitosis and mitotic spindle function were re-induced in S and/or G2 phase in drug, but the vast majority are not, showing that cells which progress through the cell cycle in drug do so without inducing many genes involved in normal growth. Conversely, amongst the 1,978 genes upregulated at Day 28 only the Pregnancy-specific beta-1-glycoproteins reached significant enrichment; upregulation of these genes is associated with aggressive tumours (52,53), but not so far with osimertinib resistance.

These results show that on entering the cell cycle, cells on drug progress through S, G2 and M, but often lose viability. Many genes required for critical biosynthetic pathways remain down-regulated in these cells, which likely contributes to the failure of the population to proliferate despite frequent entry to S phase.

### Aberrant replication fork progression and chromosome segregation in osimertinib-treated cells

*CCNA1* expression is linked to DNA damage and we hypothesised that this may result from DNA replication in drug being aberrant or stressed. We therefore applied TrAEL-seq to control and osimertinib-treated cells, which profiles the average directionality of forks across the genome to reveal the replication programme (38). TrAEL-seq is well-suited to studying drug-treated populations as no synchronisation or nucleotide analogue incorporation is required (37). We acquired two biological replicate populations at Day 0, 7, 14 and 28 of osimertinib treatment, which were processed as two multiplexed pools allowing quantitative comparison of signals across the time courses. In keeping with the EdU data, total TrAEL-seq read count decreased from Day 0 to Day 7 and never exceeded the day 0 signal although there was considerable variation in read count between the two time courses (Figure S3A).

TrAEL-seq reveals the average direction in which replication forks travel at each genomic region across a population of cells. This is well-defined in early replicating genomic regions where forks move out from Initiation Zones such that the average direction of fork movement is highly polarised (Figure 3A, Day 0, orange arrows). However, as drug treatment progresses this polarisation is lost, indicating that Initiation Zones are no longer used universally across the population and replication patterns are more random (Figure 3A, Day 7-28). This effect is prominent on a metaplot of fork directionality around all PC9 initiation zones (Figure 3B), with Day 7 and Day 14 samples showing an intermediate phenotype. Loss of polarisation could also arise through increased non-replication-linked TrAEL-seq reads obscuring the replication signal, for example from DNA breaks, but the signal is decreased rather than increased relative to Day 0 (Figure S3A), and therefore a loss of uniformity of the replication programme is the most likely explanation. In keeping with this, the decrease in TrAEL-seq read count with increasing distance from Initiation Zones, which arises from forks accelerating with distance (37,54–56), is muted in drug-treated cells, indicating less coherent use of individual Initiation Zones (Figure S3B).

**Figure 3:**
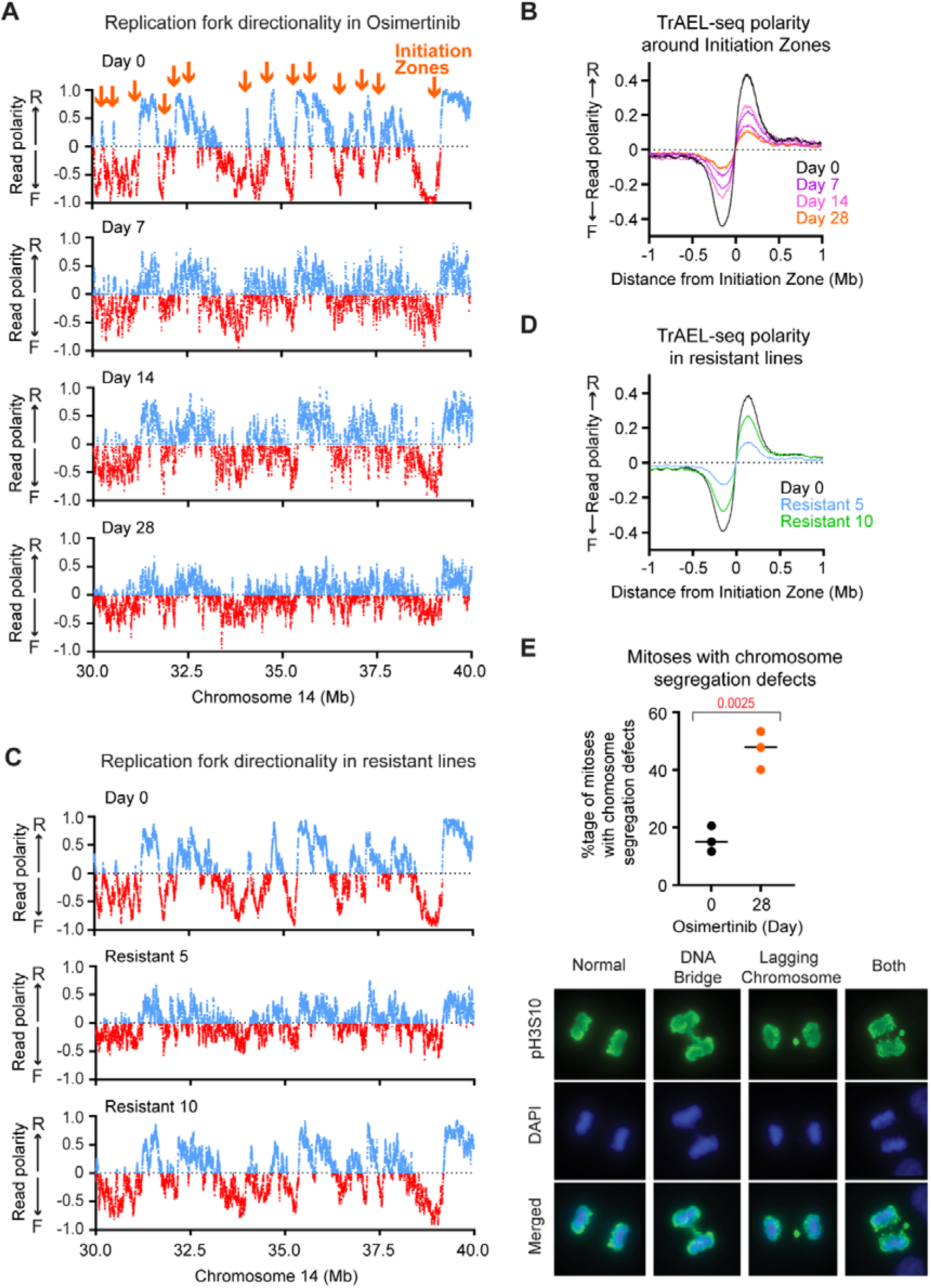
Aberrant replication fork progression and chromosome segregation in osimertinib-treated cells. A) DNA replication fork directionality across a representative 10 Mb region of chromosome 14 using TrAEL-seq in PC9 cells following 0, 7, 28, and 58 days of 500 nM osimertinib treatment. Data is an average of two biological replicates, strong Initiation Zones are highlighted in orange. Read polarity calculated as (R-F)/(R+F) in 30 kb windows spaced every 1 kb provides replication Fork Directionality (RFD). B) Metaplot of DNA replication fork directionality measured in 50 kb windows within ±1 Mb of replication initiation zones in PC9 cells following 0, 7, 14, and 28 days of 500 nM osimertinib treatment. C) Quantification of DNA replication fork directionality across a 10 Mb region of chromosome 14 in untreated and two osimertinib-resistant PC9 populations, analysed as in A. D) Metaplot of DNA replication fork directionality measured in 50 kb windows within ±1 Mb of replication initiation zones in untreated and osimertinib-resistant PC9 populations. E) Quantification of the proportion of mitoses exhibiting chromosome segregation defects, with representative images, at days 0–28 of 500 nM osimertinib treatment. Statistical significance was determined by unpaired t-test, n=3.

Strong replication Initiation Zones are associated with regions of open chromatin lying upstream or downstream of highly transcribed genes, and we asked whether gene up- and down-regulation caused by osimertinib differentially impacted Initiation Zone usage. At Day 0, TrAEL-seq polarity changes in the 50kb upstream of genes that are down-regulated by osimertinib show that Initiation Zones are concentrated in this region (Figure S3C, analysed as in (57)). This polarity change is almost absent at Day 28, showing loss of Initiation Zone activity. The equivalent analysis of the 50 kb upstream of genes that are up-regulated in osimertinib also indicates that Initiation Zones are present, but these also disappear in osimertinib (Figure S3D), so the changes in gene expression are not directly responsible for the difference in DNA replication profile.

Notably, TrAEL-seq of two osimertinib resistant lines that emerged from the DTP state did not show a complete reversion to the replication pattern of untreated cells (Figure 3C), even though these cells proliferate effectively in drug (Figure S3C). Metaplot analysis across all Initiation Zones reveals that the restoration is partial in line R10 and very low in line R5 (Figure 3D). The loss of uniformity in DNA replication pattern is therefore not simply a result of the DTP state or the reduction in S phase cells, but rather an effect of osimertinib treatment that is separable from proliferation.

Defects in DNA replication frequently result in chromosome segregation defects due to cells entering mitosis prior to completion of replication. We therefore examined Day 0 and Day 28 mitosis events for DNA bridges and lagging chromosomes, and observed that aberrant chromosome segregation occurred 3-fold more often in osimertinib-treated cells, with almost half of mitosis events being scored as aberrant (Figures 3E,F).

Therefore, replication is frequent in PC9 cells treated with osimertinib but the replication programme is profoundly disrupted, and chromosome segregation errors occur frequently.

### Increased DNA damage and repair in osimertinib-treated cells

Replication defects can lead to incomplete DNA replication and DNA damage marked by □H2A.X foci. As predicted, osimertinib treatment increases the average number of □H2A.X foci per cell (Figure 4A), with the difference driven by a greater number of cells with many foci (>20) per cell (Figure 4B). This increase in □H2A.X is suppressed by co-treatment with palbociclib (Figure 4A,B), linking DNA damage to DNA replication in osimertinib.

**Figure 4:**
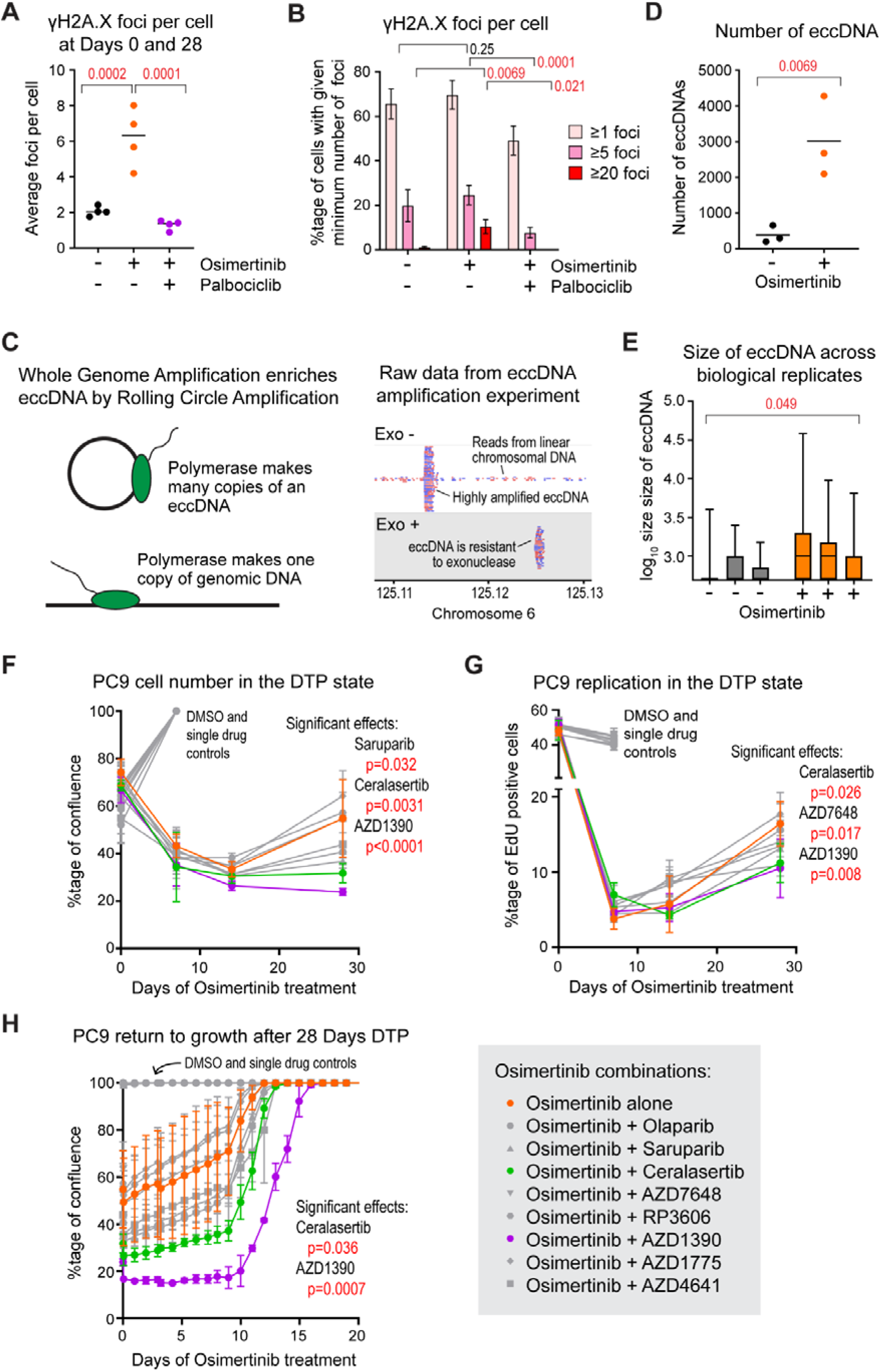
Increased DNA damage and repair in osimertinib-treated cells. A) Quantification of the proportion of cells with 1–4, 5–19, or ≥20 γH2AX foci per cell following 0 or 28 days of 500 nM osimertinib treatment ±16 nM palbociclib. Statistical significance was determined by lognormal one-way ANOVA with Sidak’s multiple comparisons correction, n=4. B) Quantification of the mean number of γH2AX foci per cell following 0 or 28 days of 500 nM osimertinib treatment ±16 nM palbociclib. Statistical significance was determined by two-way ANOVA with Tukey’s multiple comparisons correction, n=4, error bars ±1 SD. C) Schematic of the Circle-seq method, involving optional exonuclease-mediated degradation of linear DNA followed by rolling-circle amplification, resulting in sequencing libraries enriched for exonuclease-resistant circular DNA. D) Quantification of eccDNA abundance in 10,000 PC9 cells following 0 or 28 days of 500 nM osimertinib treatment. Statistical significance was determined by unpaired t-test on log-transformed data, n=3. E) Quantification of eccDNA size distributions across three biological replicates of 10,000 PC9 cells following 0 or 28 days of 500 nM osimertinib treatment. Statistical significance was determined by nested t-test with Dunn’s multiple comparisons correction, n=3. F) Quantification of PC9 cell confluence over 0, 7, 14, and 28 days of treatment with DMSO control, single agents, or combination treatments with osimertinib. Statistical significance was determined by two-way ANOVA comparing combination treatments to single-agent osimertinib, with Dunnett’s correction, n=3, error bars ±1 SD. G) Quantification of the proportion of PC9 cells positive for EdU over 0, 7, 14, and 28 days of treatment with DMSO control, single agents, or combination treatments with osimertinib. Statistical significance was determined by two-way ANOVA with Dunnett’s correction, n=3, error bars ±1 SD. H) Quantification of time to regrowth (to 100% confluence) following drug withdrawal after 28 days of treatment with DMSO control, single agents, or combination treatments with osimertinib. Statistical significance was determined by one-way ANOVA on area under the curve (AUC) values with Dunnett’s multiple comparisons correction, n=3, error bars ±1 SD.

Resolving individual damage or repair products in chromosomal DNA is extremely challenging, but DNA repair events are known to form extrachromosomal DNA circles (eccDNA) (58–60), which can be detected at single molecule sensitivity by rolling circle amplification (61–63). eccDNA are abundant and heterogenous, so to resolve individual species we optimised an assay for samples of 1,000 cells, and included an exonuclease treatment to which eccDNA are resistant. Strong signals from ∼1-50kb regions were observed, and these were resistant to exonuclease treatment showing them to be eccDNA (Figure 4C). Rolling circle amplification is sensitive but poorly quantitative, and we noted that exonuclease treated samples reported very variable eccDNA abundance. However, in samples without exonuclease digestion the rolling circle amplification competes with whole genome amplification of the much more abundant genomic DNA, which limited the amplification and yielded more reproducible results. Using this method we determined that after 28 days of osimertinib treatment, the number of eccDNA detected per 1,000 cells rose ∼5-fold and the number of larger circles (>2k) rose ∼200-fold (Figure 4D,E), consistent with increased DNA repair.

Given this evidence for DNA damage and repair in the DTP state, we tested eight DNA damage response inhibitors to determine whether inhibiting repair pathways can act synergistically with osimertinib to prevent survival of the DTP population. The set included ATM (AZD1390), ATR (ceralasertib) and DNA-PKcs inhibitors (AZD7648) to impair damage signalling, PARP (Olaparib, Saruparib) and POLθ inhibitors (AZD4641) to block break repair, and MYT1 (RP3606) and WEE1 (AZD1775) inhibitors to modulate entry to mitosis in the presence of DNA damage. Assays were performed in PC9 and HCC4006 cells, with each drug at a concentration that did not impair growth as a single agent based on previous studies or growth response curves in these cell lines (Figure S4A). We measured cell number as percentage confluency over 28 days for PC9 and 14 days for HCC4006, percentage EdU positive cells, and rate of return to growth on drug removal at various times (Figure 4F-H, S4B-D), though measurements of single drug controls were stopped on reaching confluence. Results were highly consistent across assays and cell lines, with AZD1390 and ceralasertib inhibitors reducing cell number, replication and return to growth in combination with osimertinib showing that effective DNA damage responses are critical for survival in the DTP state.

The DNA replication defects in osimertinib-treated cells therefore have direct consequences for genome stability, resulting in DNA damage and requiring ongoing DNA repair to maintain cell viability.

## Discussion

Our work reveals that the DTP state can be far more dynamic than previously thought, with cells rapidly dividing and dying at similar rates such that the population remains approximately constant. However, the cell cycle occurs with improper gene expression, defective DNA replication, and increased DNA damage.

DNA replication in the DTP state does not result from a subpopulation of pre-existing resistant cells, and instead represents stochastic escape from arrest at the G1/S checkpoint, with reinforcement of the checkpoint by palbociclib dramatically decreasing escape. We have observed this phenotype in multiple cell types with both EGFR and MEK inhibitors (28), indicating that cell cycle arrest is incomplete in cells which survive loss of growth signalling through the MEK-ERK pathway. However, loss of MEK-ERK signalling does suppress gene expression programmes required for growth, such as ribosome synthesis, which we suggest underlies arrest and cell death during cell cycles in the DTP state, in turn preventing population expansion despite ongoing cell cycles. Importantly, the ongoing proliferation acts to maintain the DTP state as loss of proliferation in osimertinib + Palbociclib leads to progressive loss of cells, meaning that the DTP state also involves gradual cell death unlinked to proliferation.

Once cells enter S phase in presence of osimertinib, the cell cycle is profoundly abnormal based on TrAEL-seq, segregation analysis, □HA.X staining and susceptibility to ATM and ATR inhibitors. This provides an explanation for the previously reported induction of DSBs by oncogene-targeted therapies and susceptibility of treated cells to ATM inhibition (64). Major DNA replication and repair proteins are induced on entry to S phase in drug, and the subset of replication factors we previously noted not to be induced under Selumetinib treatment are induced in osimertinib, so we cannot ascribe the replication defects and DNA damage to loss of expression of any individual gene(s). More likely, these arise from cells entering the cell cycle without the appropriate preparation in G1 to ensure the supply of metabolites and biosynthetic capability.

Perhaps unsurprisingly, the replication profile is also unusual in osimertinib. EdU incorporation assays show that the amount of DNA replication per cell is normal, but TRAEL-seq indicates that canonical initiation zones are not used as frequently. In other words, the same amount of replication forks must be active but they do not initiate in the same places. Single molecule analysis of DNA replication has revealed frequent origin firing outside canonical Initiation Zones (65), so presumably under osimertinib treatment the balance moves more towards stochastic origin firing and away from use of the defined Initiation Zones. This could be due to DNA damage disrupting the distribution of replication timing factor Rif1, which is critical for defining Initiation Zones (66,67), but could be an indirect effect of gene expression changes since origin firing often occurs at the 5’ end of highly expressed genes (68,69). Both up- and down-regulated genes lose Initiation Zone activity by Day 28, but our designation of up and down-regulation is based on RNA-seq normalised to average gene expression; any global reduction in transcription, which is very likely under long-term EGFR inhibition, will not be detected. Such global effects will need to be probed by direct measurements of transcription and chromatin accessibility.

The emergence of resistance from the DTP state provides a model for the known emergence of resistance in patients (70–74). DTP cells can revert to a proliferative state on drug removal by non-genetic mechanisms, but drug-resistant cells also eventually emerge from the DTP state under continued drug treatment (19,26). Here, we have addressed this emergence of genetic resistance: we have previously hypothesised that inappropriate DNA replication in the presence when the MEK/ERK pathway is inhibited can promote DNA damage and mutation (28), and here we provide multiple lines of evidence that this is the case for cells treated with the EGFR inhibitor osimertinib. Altogether this leads to a model in which the clinical utility of monotherapies which target MEK/ERK or upstream growth signalling is limited due to inherently pro-mutagenic replication when cells escape, leading to a spectrum of *de novo* resistance. However, this also represents a vulnerability that may be exploited by combination therapies, using DDR inhibitors in combination with growth inhibitors to specifically target cells that would otherwise go on to develop resistance.

## Acknowledgements

We greatly appreciate the assistance of Simon Walker, Hanneke Okkenhaug, Sam Thompson, Yulia Chupalova, Kleopatra Dagla, Rita Dapaah, Simon Andrews, Sarah Inglesfield and Laura Biggins in conducting these experiments. TrAEL-seq, DNA-seq and RNA-seq library sequencing and processing were performed by the Genomics (Geno06) and Bioinformatics (Bioinf01) Facilites, flow cytometry and imaging were performed in the Flow Cytometry (Flowcyt04) and Imaging (Imag07) Facilities, which receive financial support from the Institute Core Capability Grant.

---

**Supplementary Figure 1:**
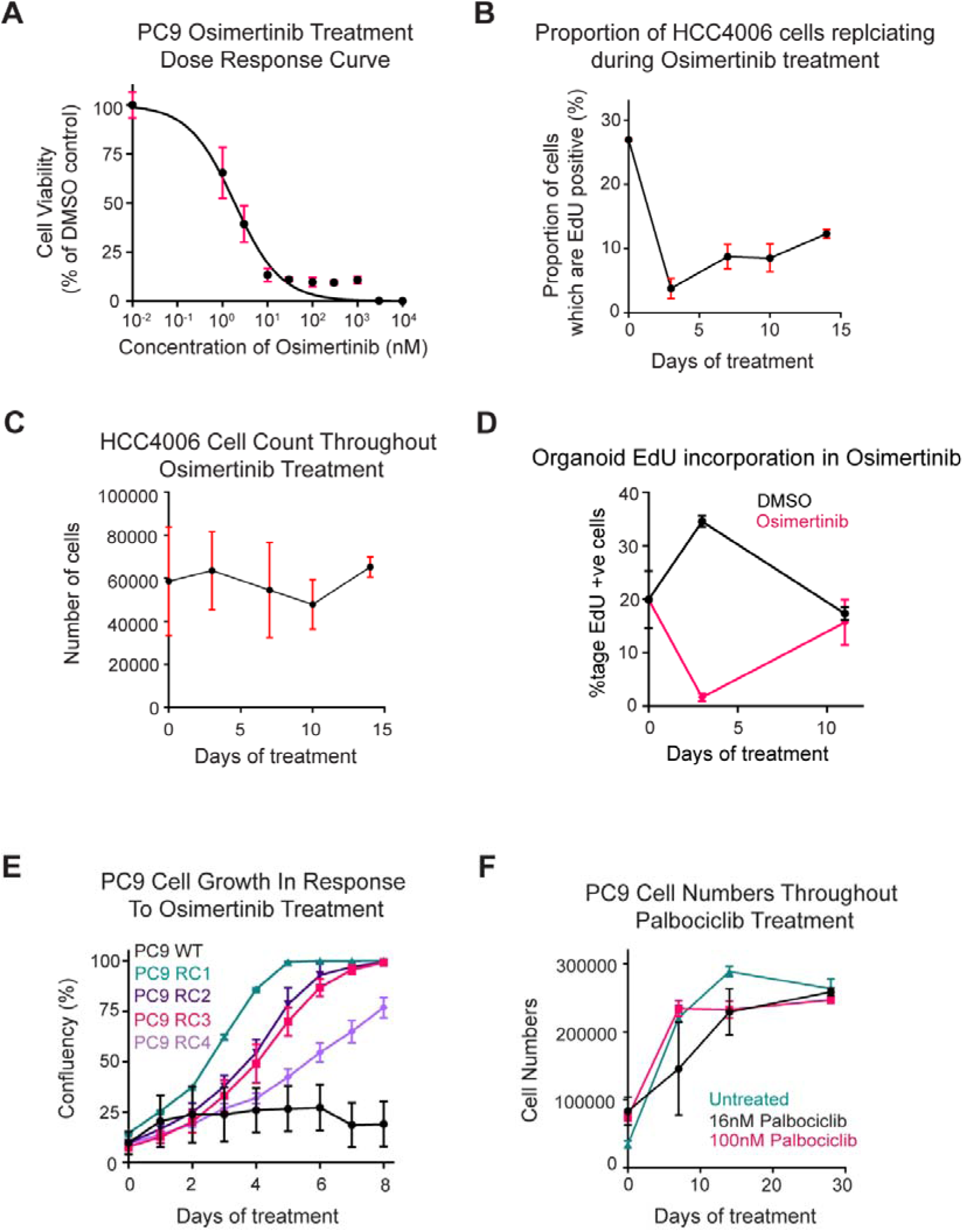
Supplement to Extensive DNA replication occurs in the DTP state. A) Dose–response curve showing viability of PC9 cells following 4 days of osimertinib treatment (0.01 nM to 10 µM, half-log scale), normalised to matched DMSO controls. Curves were fitted using non-linear regression (log[inhibitor] vs. normalised response, variable slope), error bars ±1 SD. B) Quantification of the proportion of HCC4006 cells positive for EdU over 14 days of 500 nM osimertinib treatment, n=3, error bars ±1 SD. C) Quantification of HCC4006 cell numbers over 14 days of 500 nM osimertinib treatment, n=3, error bars ±1 SD. D) Quantification of the proportion of NSCLC organoid cells (Tempus Labs, harbouring a heterozygous L858R EGFR mutation) positive for EdU over 14 days of 500 nM osimertinib treatment, n=3, error bars ±1 SD. E) Quantitation of the time required for wild-type PC9 and multiple osimertinib-resistant PC9 clones take to reach full confluence under continuous 500nM osimertinib treatment, n=3, error bars ±1 SD. F) Quantification of PC9 cell numbers over 28 days of treatment with 0, 16 or 100 nM Palbociclib, n=3, error bars ±1 SD.

**Supplementary Figure 2:**
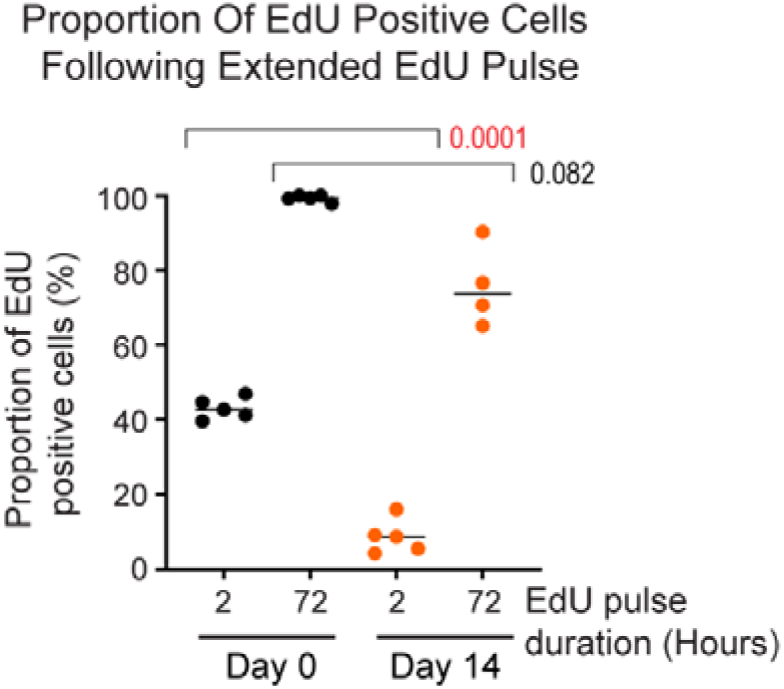
Supplement to the DTP cell cycle involves frequent cell death and aberrant gene expression. Quantitation of the proportion of PC9 cells positive for EdU incorporation following 2-hour or 72-hour exposure to 0.5 µM EdU after 0 or 14 days of 500 nM osimertinib treatment. Significance was determined by one way ANOVA, n=4-5.

**Supplementary Figure 3:**
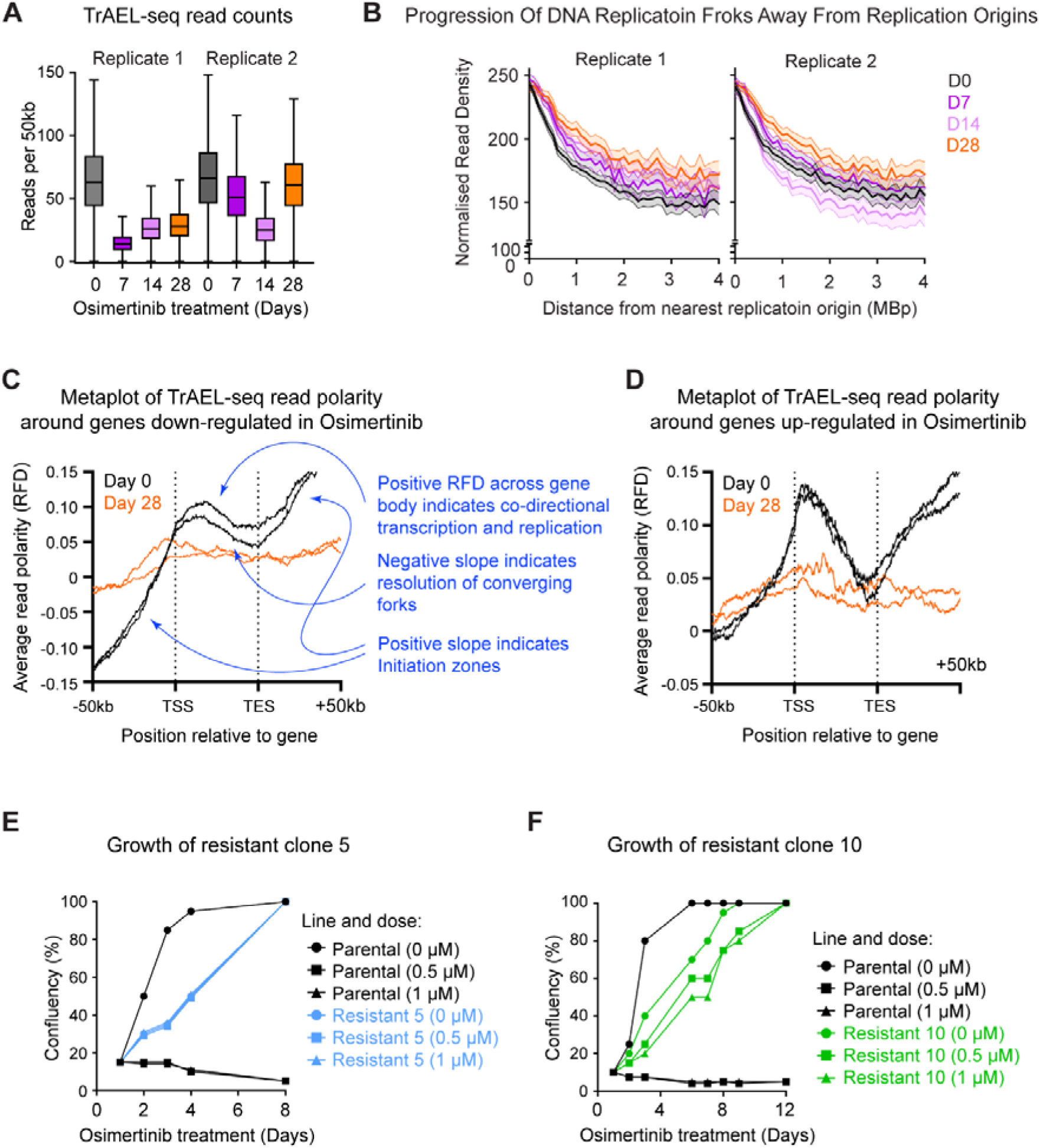
Supplement to aberrant replication fork progression and chromosome segregation in osimertinib-treated cells. A) Quantitation of TrAEL-seq read density in PC9 following 0, 7, 14, or 28 days of 500 nM osimertinib treatment. B) Quantitation of normalised TrAEL-seq read density in 50 Kb windows as a function of distance from the nearest DNA replication origin. Solid lines represent the mean, with shaded regions indicating standard deviation. C) Quantitation of average TrAEL-seq signal across genes down-regulated following osimertinib treatment relative to untreated cells, including +/− 50 Kbp flanking regions. D) Quantitation of average TrAEL-seq signal across genes up-regulated following osimertinib treatment relative to untreated cells, including +/− 50 Kbp flanking regions. E) Growth curves of parental PC9 cells and osimertinib resistant clone 5 treated with 0, 0.5 or 1 µM osimertinib over 8 days. F) Growth curves of parental PC9 cells and osimertinib resistant clone 10 treated with 0, 0.5 or 1 µM osimertinib over 8 days.

**Supplementary Figure 4:**
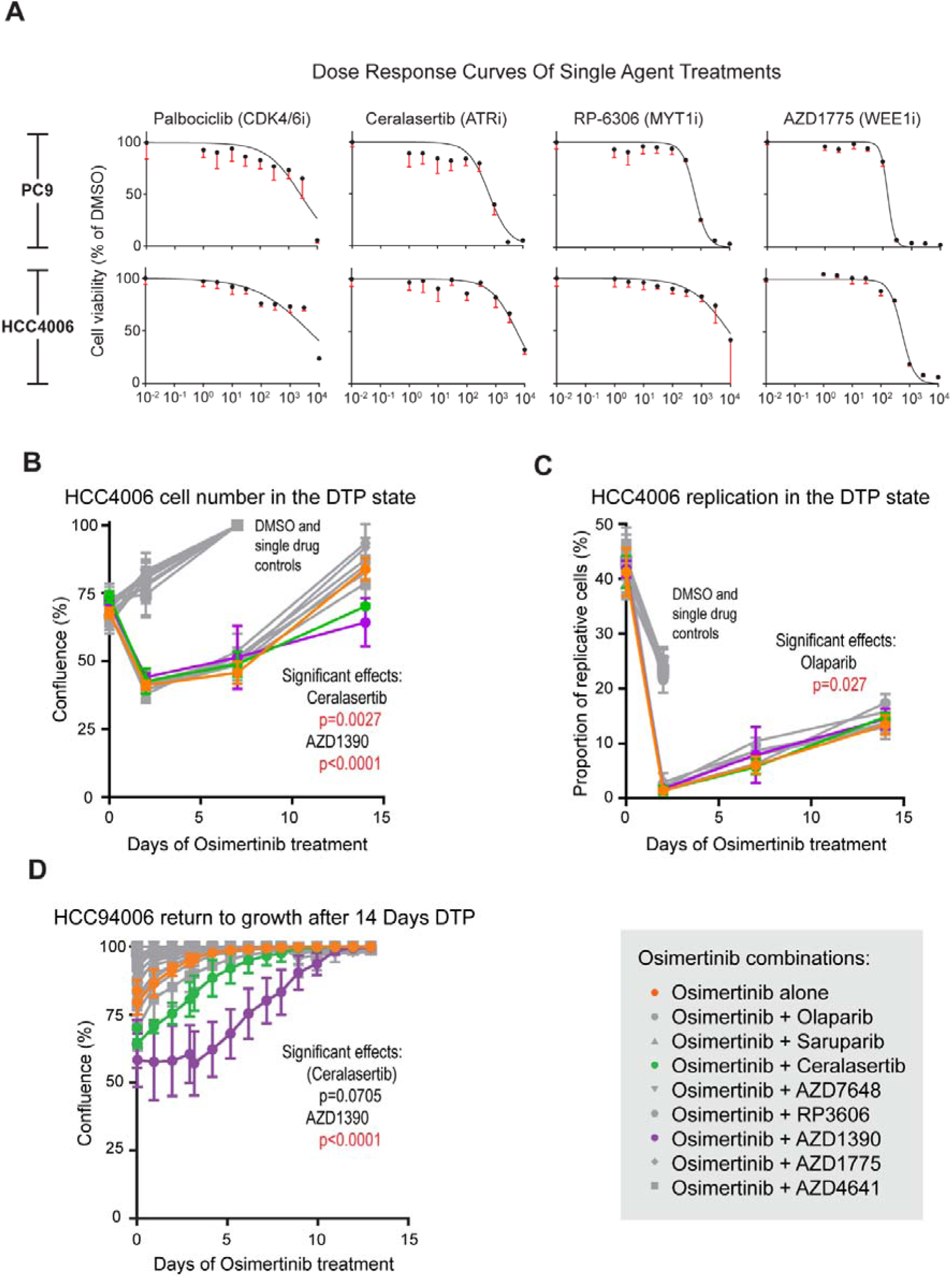
Supplement to Increased DNA damage and repair in osimertinib-treated cells. A) Dose–response curves showing viability of PC9 (top) and HCC4006 (bottom) cells following 4 days of treatment with Palbociclib (CDK4/6 inhibitor), Ceralasertib (ATR inhibitor), RP-6306 (MYT1 inhibitor), or AZD1775 (WEE1 inhibitor) (0.01 nM to 10 µM, half-log scale), normalised to matched DMSO controls. Curves were fitted using non-linear regression (log[inhibitor] vs. normalised response, variable slope), n=3, error bars ±1 SD. B) Quantification of HCC4006 cell confluence over 0, 3, 7, and 14 days of treatment with DMSO control, single agents, or combination treatments with osimertinib, n=3, error bars ±1 SD. C) Quantification of the proportion of HCC4006 cells positive for EdU over 0, 3, 7, and 14 days of treatment with DMSO control, single agents, or combination treatments with osimertinib, n=3, error bars ±1 SD. D) Quantification of time to regrowth (to 100% confluence) of HCC4006 cells following drug withdrawal after 14 days of treatment with DMSO control, single agents, or combination treatments with osimertinib, n=3, error bars ±1 SD.

